# A multiwell-plate *Caenorhabditis elegans* assay for assessing the therapeutic potential of Bacteriophages against Clinical Pathogens

**DOI:** 10.1101/2022.01.05.474866

**Authors:** Prasanth Manohar, Belinda Loh, Namasivayam Elangovan, Archana Loganathan, Ramesh Nachimuthu, Sebastian Leptihn

## Abstract

In order to establish phage therapy as a standard clinical treatment for bacterial infections, testing of every phage to ensure the suitability and safety of the biological compound is required. While some issues have been addressed over recent years, standard and easy-to-use animal models to test phages are still rare. Testing of phages in highly suitable mammalian models such as mice is subjected to strict ethical regulations, while insect larvae such as the *Galleria mellonella* model suffers from batch-to-batch variations and requires manual operator skills to inject bacteria, resulting in unreliable experimental outcomes. A much simpler model is the nematode *Caenorhabditis elegans* which feeds on bacteria, a fast growing and easy to handle organism which can be used in high-throughput screening. In this study, two clinical bacterial strains of *Escherichia coli*, one *Klebsiella pneumoniae* and one *Enterobacter cloacae* strain were tested on the model system together with lytic bacteriophages that we isolated previously. We developed a liquid-based assay, in which the efficiency of phage treatment was evaluated using a scoring system based on microscopy and counting of the nematodes, allowing increasing statistical significance compared to other assays such as larvae or mice. Our work demonstrates the potential to use *Caenorhabditis elegans* to test the virulence of strains of *Klebsiella pneumoniae, Enterobacter cloacae* and EHEC/ EPEC as well as the efficacy of bacteriophages to treat or prevent infections, allowing a more reliable evaluation for the clinical therapeutic potential of lytic phages.

**Importance:** Validating the efficacy and safety of phages prior to clinical application is crucial to see phage therapy in practice. Current animal models include mice and insect larvae, which pose ethical or technical challenges. This study examined the use of the nematode model organism, *C. elegans* as a quick, reliable and simple alternative for testing phages. The data shows that all the four tested bacteriophages can eliminate bacterial pathogens and protect the nematode from infections. Survival rates of the nematodes increased from <20% in the infection group to >90% in the phage treatment group. Even the nematodes with poly-microbial infections recovered during phage cocktail treatment. The use of *C. elegans* as a simple whole-animal infection model is a rapid and robust way to study the efficacy of phages before testing them on more complex model animals such as mice.

## Introduction

Gram-negative bacteria can cause serious infections in humans, including strains from the family of the *Enterobacteriaceae* (*Escherichia coli*, *Klebsiella pneumoniae* and *Enterobacter cloacae*). Over the years, such pathogenic strains have been exhibiting increasing resistance to last-resort antibiotics such as carbapenems and colistin (1,2,3). With the decreasing number of available antibiotics, global overuse and misuse in both clinical medicine and agriculture, antimicrobial resistance is an undeniable reality which is also unavoidable (4). However, with appropriate governmental regulations concurrent with the education of doctors and patients alike, the velocity at which resistant strains emerge may be decreased. Nonetheless, novel antimicrobial compounds that can be used in medicine and/or agriculture would not only benefit the sectors but are essential for global health. In light of the withdrawal of pharmaceutical global players to discover new classes of small molecule antibiotics, an alternative approach is to use bacteriophages as biological therapeutics (5,6).

Phage therapy is the application of bacterial viruses that kill its host and thus eliminate a bacterial infection. Although phage therapy is an old concept, it is necessary to improve the understanding of antibacterial properties of phages and their applications before bringing phages into clinical practice (7). Bacteriophages are known to be genus-, species-, or even strain-specific and exhibit no or only minimal side effects when infecting pathogens inside a mammalian host (8,9,10). Their interactions within mammalian hosts, in particular with the immune system, are complex and a healthy phageome has been shown to be beneficial to the host (11,12,13,14). In order to translate a phage isolate into a “drug”, several challenges need to be addressed, such as evaluating host susceptibility to the phage to ensure that the phage can infect and kill the pathogen, genomic analysis to guarantee the absence of virulence factors or lysogeny genes, producing phages at high numbers devoid of any toxic compounds such as bacterial LPS, and testing the efficacy of the phage in an animal model (15,16).

There is a need to improve the current work flow - from its discovery to the application of phages, which includes investigating the efficacy of a phage in a simple animal model (17,18). Many studies have reported the use of waxworms (*Galleria mellonella*) as a model to study the efficacy of bacteriophages (15,19,20,21). However, it requires a skilful operator to inject bacteria and / or phages into individual larvae which sometimes could result in large variations between batches, and thus lead to non-reproducible observations. While ethical approval is easier to obtain for work with *G. mellonella*, large numbers of larvae are needed to achieve statistical relevance and therefore a considerable amount of time is required to reliably test a phage. While a mammalian host such as mice would be most suitable to test a phage prior to clinical use, ethical approval is difficult to obtain, and experiments take too long (especially for emergency use of phages in compassionate therapy), while low animal numbers also decrease statistical significance. Thus, a robust, reliable and reproducible assay with large numbers of animals would be highly advantageous to test the efficacy of a phage. In this study, we used the nematode *Caenorhabditis elegans* as a simple whole-animal infection model and established a liquid-based assay in a multi-well format. *C. elegans* is an established animal infection model to study pathogenesis and to evaluate the efficacy of drugs (22,23,24,25,26). However, and perhaps surprisingly, there are very limited studies that report the use of *C. elegans* as an animal model to evaluate the efficacy of phage therapy (27,28). Studies have been conducted to investigate virulence mechanisms of *Pseudomonas aeruginosa* (29), *Burkholderia paseudomallei* (30) and *Salmonella typhimurium, S. enteritidis* and *S. pullorum* (31,32,33). In addition, infections of *C. elegans* by pathogenic *E. coli*, *K. pneumoniae*, *E. cloacae* and the Gram-positive *Staphylococcus aureus* have been studied where the mechanisms of host-pathogen interaction were elucidated, but also the antibacterial activity of phages were examined (22,27). In all of these studies, solid media was used in the experiments, making it a complex, and possibly sub-optimal assay. The goal of our study was to establish a *C. elegans* – pathogen (*E. coli*, *K. pneumoniae* and *E. cloacae*) liquid-based platform to elucidate the efficacy of lytic phages *in vivo*. Here, we were able to demonstrate that in the presence of lytic phages the lifespan of infected nematodes was increased up to 6-fold compared to the controls without phage. In the bacteria infected groups, the nematode survival was reduced by ∼80% within 4 days. Interestingly, the prophylactic treatment (phages introduced one hour before bacterial infection) showed better efficacy than the therapeutic treatment group (phages introduced 2 hours after bacterial infection) across all four tested pathogens.

## Results

### C. elegans as infection models

We established a *C. elegans* liquid-based assay with the aim of providing a robust and reliable approach with high numbers of repeats for statistical significance, to study the virulence of bacterial isolates *in vivo* and the effects of bacteriophages that target these pathogens. Prior to experimental conditions, we conducted a series of control experiments. In our first control, *C. elegans* were fed with the non-pathogenic *E. coli* strain OP50 and were observed to live up to 5 days. In the case where no bacteria were added as feed, the lifespan of *C. elegans* was not drastically reduced despite the starvation conditions (S.fig.1).

It had previously been shown that several pathogens have the ability to kill *C. elegans* including *Pseudomonas and Klebsiella* when the nematodes were exposed to the bacteria in the surrounding media. Thus, nematodes can be used as a model organism to study bacterial infection (34,35). In addition, despite being the bacterium that *C. elegans* can feed on, studies have shown that certain *E. coli* strains which are pathogenic in humans or animals have the ability to infect nematodes as well (36). Therefore, in our second set of control experiments, we sought to determine the pathogenicity of bacterial strains on *C. elegans*. Groups of 20 nematodes were independently infected with 4 pathogenic strains, *Escherichia coli* 131, *Escherichia coli* 311, *Klebsiella pneumoniae* 235 and *Enterobacter cloacae* 140 and observed over 5 days. Regardless of strain, when the nematodes were infected with 10^3^ CFU/mL of bacterial pathogen, they lived beyond five days whereas 10^7^ CFU/mL caused death within 3 days (S.fig.2).

As our experimental set-up was conducted in 96-well plates, the maximum volume in each well was 100 µL. Over extended durations lasting longer than 5 days, the experimental conditions would lead to volume reduction before eventually drying up completely, thus altering the conditions and potentially affecting the results of our study. Therefore, in this study, a bacterial concentration of 10^5^ CFU/mL was chosen as all nematodes succumbed to infection in less than 5 days.

Nematodes were evaluated based on appearance (reflectance and optical transparency) and mobility (alive: moving; dead: inactive) as observed under an inverted microscope (40x magnification). A representative scoring model is provided in Fig.1. Scoring was performed in 24 hour intervals. We classified phages to be effective if nematode survival was higher than 70% of the infected nematodes while less than 20% survival in the infected groups were classified as ineffective.

**Figure 1:**
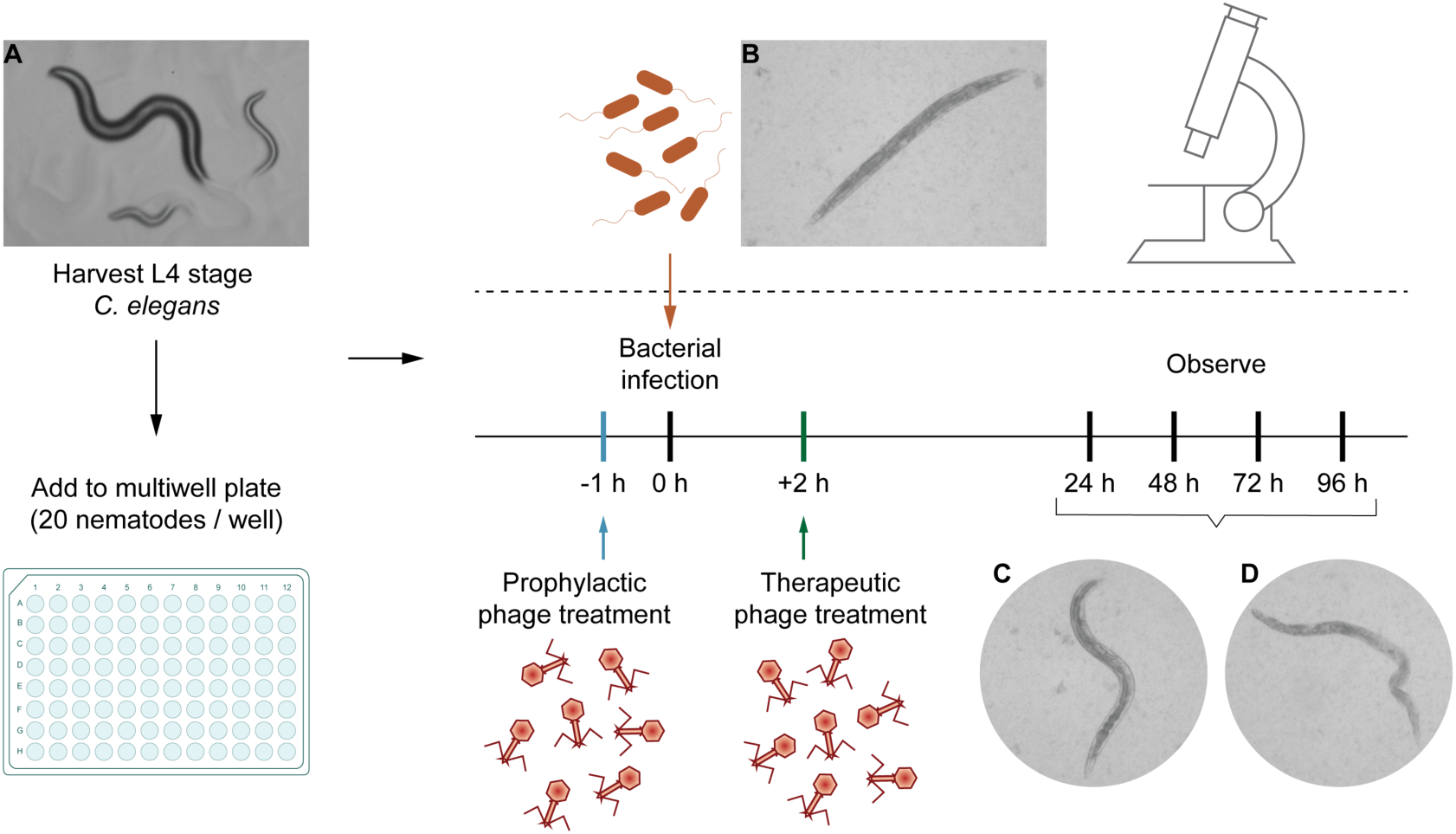
Schematic representation of the assay and distinct appearances of the nematode *C. elegans* on solid or in liquid medium. Representative microscopic images may serve as a guide for scoring by the operator. Movement is an important indicator for the viability of a worm but not a strict requirement to score a nematode to be alive. (A) *C. elegans* on NGA plates, (B, C) Live L4 stage mature nematodes in liquid medium. (D) Dead nematode after bacterial infection. The worms were examined under an inverted microscope at 40x magnification.

In order to determine if our model organism is sensitive to endotoxin, four live or heat-inactivated pathogenic bacterial strains (*E. coli* 131, *E. coli* 311, *K. pneumoniae* 235 and *E. cloacae* 140) were tested on the nematodes. When the *C. elegans* were infected with live pathogenic *E. coli*, *K. pneumoniae* or *E. cloacae* strain, a significant reduction in nematode survival was observed. In contrast, nematode survival was maintained at 95% when infected with heat-inactivated bacteria indicating that heat stable molecules such as the endotoxin LPS, do not lead to mortality of the worms (S.fig.3). Thus, our experiments show that it is indeed the pathogenicity of the bacteria used in this study that kill the nematode.

### Phage therapy enhanced the survival of infected nematodes

The first isolate we tested in our study was *E. coli* 131, a clinical strain from a diagnostic centre in Chennai (India), isolated from blood and was previously identified to be enteropathogenic (an EPEC strain) by polymerase chain reaction (PCR) [39, 40]. We established the parameter of TD_mean_ which we defined as the mean time until we observed the nematodes to die. The TD_mean_ for *E. coli* strain 131-infected worms was found to be 3 days, a significant reduction from the usual lifespan of 5 days (Fig.2A). *Escherichia* phage myPSH1131 was previously isolated and identified to infect and lyse *E. coli* strain 131 effectively [39,40]. When myPSH1131 was tested alone, no adverse effects on *C. elegans* were observed (Fig.2A). This shows that the phage preparations were either free of toxic substances that could decrease the *C. elegans’* lifespan or the nematode is not affected by any of the components contained in the solution. In order to study the efficacy of phage treatment, varying ratios of bacteria to phage were tested i.e. 1:1, 1:10 and 1:100 which were 10^5^ CFU/mL: 10^5^ PFU/mL, 10^5^ CFU/mL: 10^6^ PFU/ mL and 10^5^ CFU/mL: 10^7^ PFU/mL. In the therapeutic treatment group, phage was added after exposure of the nematode to the pathogen. Here, the *Escherichia* phage myPSH1131 was able to increase survival up to 90% when bacteria to phage ratio was 1:100 i.e. 10^5^ CFU/mL against 10^7^ PFU/mL (Fig.2D). Other ratios tested, i.e: 1:1 and 1:10 also showed survival up to 5 days (Fig.2B,C). Similar results were observed for the prophylactic treatment group, with nematode survival up to 90% after 5 days in 1:100 (Fig.2D). In this group, the phage was added one hour before the pathogen was included into the media.

**Figure 2:**
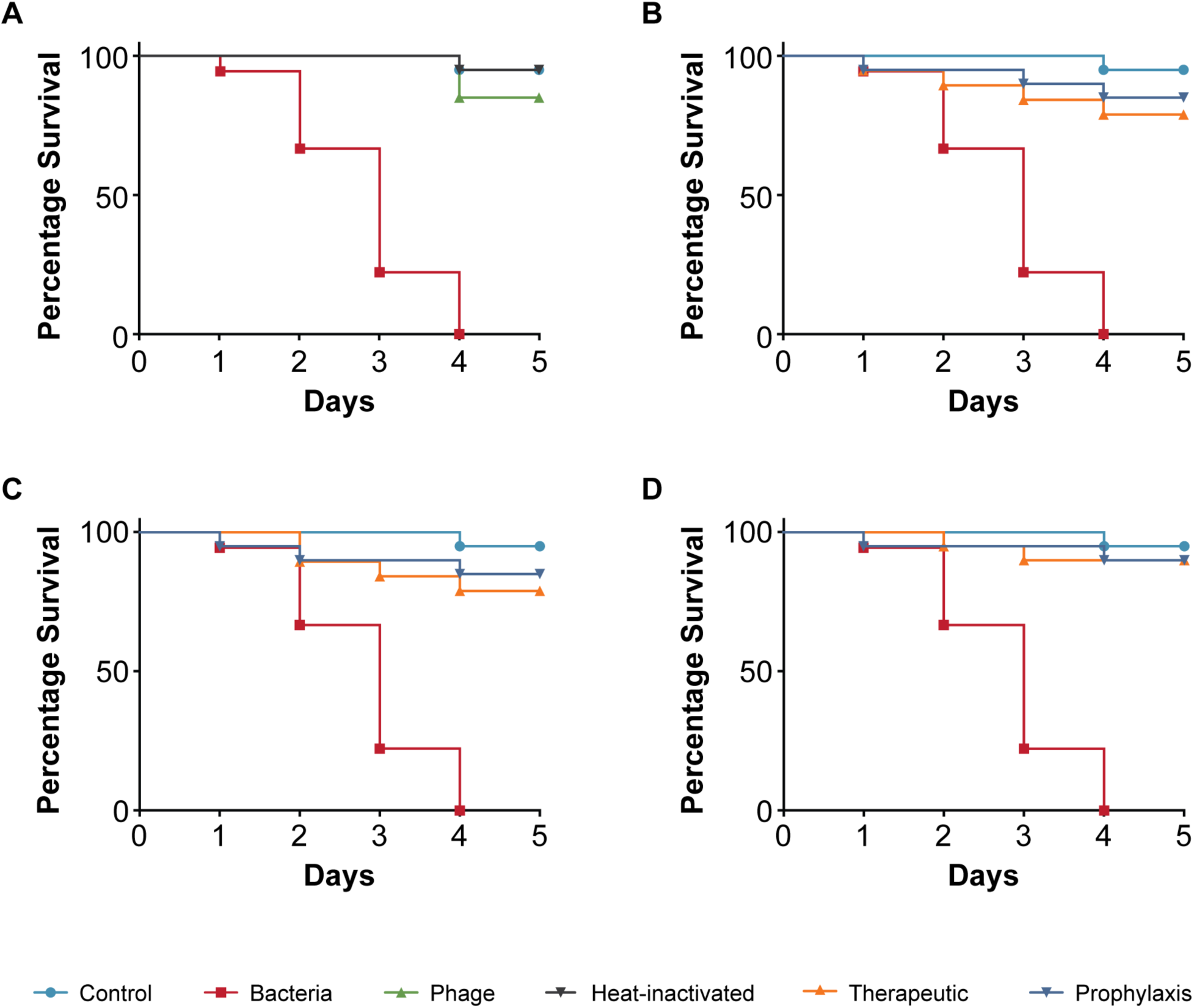
Pathogenicity of *Escherichia coli* strain 131 in *C. elegans* and efficacy of *Escherichia* phage myPSH1131 against *E. coli* infections. The control group consisted of *C. elegans* fed with *E. coli* OP50 and exposed to *E. coli* strain 131 (OD_600_=0.6) that kills *C. elegans* in liquid medium. 20 nematodes were used in each group. Representative survival curves of *C. elegans* following infection by *E. coli* strain 131 in **(A)** liquid medium consisting of M9 buffer and *E. coli* culture or *Escherichia* phage or heat-inactivated bacteria and **(B,C,D)** survival curves of *C. elegans* following infection with *E. coli* strain 131 and treatment with *Escherichia* phage, therapeutic and prophylactic treatment. **(B)** Survival curves of *C. elegans* infected and treated with bacteria and phage ratio of 1:1 i.e. 10^5^ CFU/mL and 10^5^ PFU/mL, **(C)** Survival curves of *C. elegans* infected and treated with bacteria and phage ratio of 1:10 i.e. 10^5^ CFU/mL and 10^6^ PFU/mL, **(D)** Survival curves of *C. elegans* infected and treated with bacteria and phage ratio of 1:100 i.e. 10^5^ CFU/mL and 10^7^ PFU/mL. The survival curves were plotted using Kaplan-Meier method and log-rank test was used to analyze the difference in survival rates in GraphPad Prism 7.0. A statistically significant difference (p<0.05) was observed in the treatment groups.

Next, we tested the *E. coli* strain 311 that was isolated from blood and found to be enterohemorrhagic, i.e. an EHEC strain as identified by PCR to classify pathotypes. Here, the TD_mean_ in infected *C. elegans* was 3 days and no toxicity was observed when the strain specific phage myPSH2311 was used alone (Fig.3A). *E. coli* strain 311-infected nematodes treated with the *Escherichia* phage, resulted in 90% survival after 5 days in both therapeutic and prophylactic groups (Fig.3D). At lower phage concentrations however, 1:1 and 1:10, the survival was up to 80% after 4 days (Fig.3B,C).

**Figure 3:**
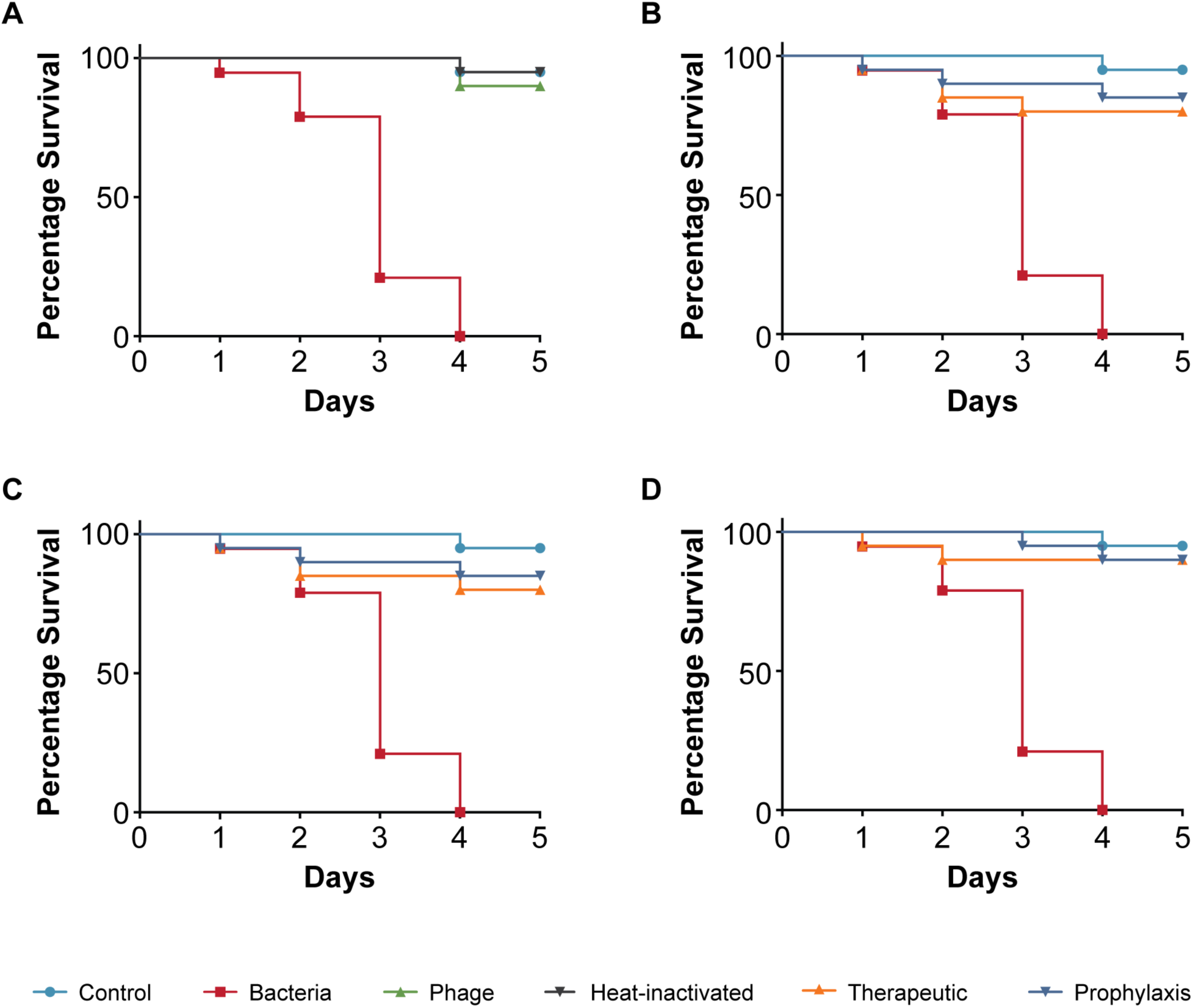
Pathogenicity of *Escherichia coli* strain 311 in *C. elegans* and efficacy of *Escherichia* phage myPSH2311 against *E. coli* infections. The control group consisted of *C. elegans* feed with *E. coli* OP50 and *E. coli* strain 311 (OD_600_=0.6) that kills *C. elegans* in liquid medium. 20 nematodes were used in each group. Representative survival curves of *C. elegans* following infection by *E. coli* strain 311 in **(A)** liquid medium consisting of M9 buffer and *E. coli* culture or *Escherichia* phage or heat-inactivated bacteria and **(B,C,D)** survival curves of *C. elegans* following infection with *E. coli* strain 311 and treatment with *Escherichia* phage, therapeutic and prophylactic treatment. **(B)** Survival curves of *C. elegans* infected and treated with bacteria and phage ratio of 1:1 i.e. 10^5^ CFU/mL and 10^5^ PFU/mL, **(C)** Survival curves of *C. elegans* infected and treated with bacteria and phage ratio of 1:10 i.e. 10^5^ CFU/mL and 10^6^ PFU/mL, **(D)** Survival curves of *C. elegans* infected and treated with bacteria and phage ratio of 1:100 i.e. 10^5^ CFU/mL and 10^7^ PFU/mL, The survival curves were plotted using Kaplan-Meier method and log-rank test was used to analyze the difference in survival rates in GraphPad Prism 7.0. A statistically significant difference (p<0.05) was observed in the treatment groups.

Next, we tested *K. pneumoniae* strain 235, which was isolated from the blood of a patient. Infection with *K. pneumoniae* 235 was significantly more virulent to *C. elegans* compared to the two *E. coli* strains we had tested. The infection killed all the nematodes within 4 days and the TD_mean_ was established to be less than 3 days (Fig.4A). The addition of phage only had no adverse effects on nematode survival. When phages were added to nematodes that had been exposed to *K. pneumoniae* strain 235, i.e. the treatment group, nematode survival increased to 80% irrespective of the phage concentrations that we analysed and the worms stayed alive at 5 days (Fig.4B,C,D). The prophylactic treatment had a slightly increased positive impact on nematode survival when bacteria to phage ratio were 1:100 with 90% survival after 5 days, compared to the therapeutic group (Fig.4D).

**Figure 4:**
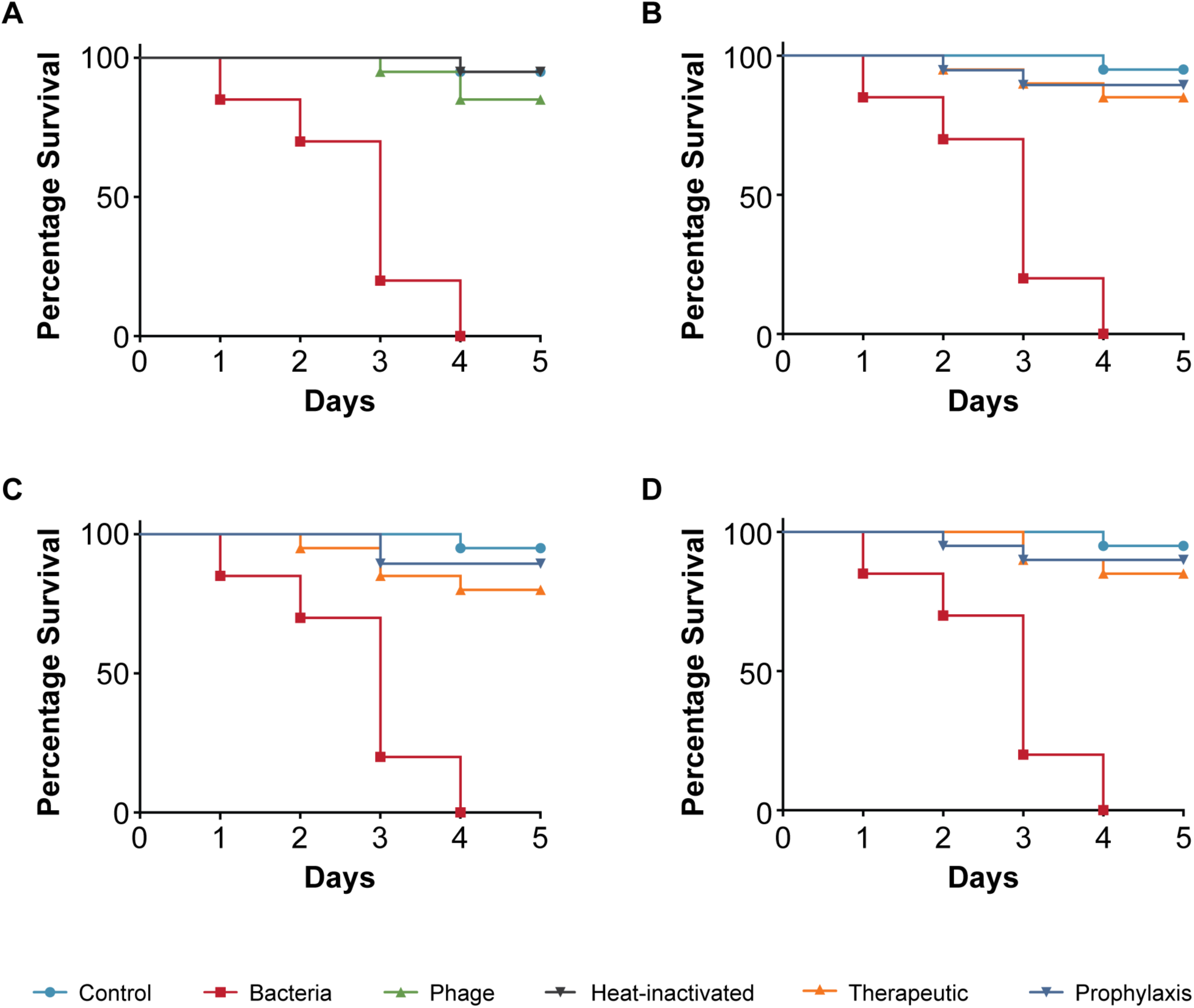
Pathogenicity of *Klebsiella pneumoniae* strain 235 in *C. elegans* and efficacy of *Klebsiella* phage myPSH1235 against *K. pneumoniae* infections. The control group consisted of *C. elegans* fed with *E. coli* OP50 and exposed to *K. pneumoniae* strain 235 (OD_600_=0.6) kills *C. elegans* in liquid medium. 20 nematodes were used in each group. Representative survival curves of *C. elegans* following infection by *K. pneumoniae* strain 235 in **(A)** liquid medium consisting of M9 buffer and *K. pneumoniae* culture or *Klebsiella* phage or heat-inactivated bacteria and **(B,C,D)** survival curves of *C. elegans* following infection with *K. pneumoniae* strain 235 and treatment with *Klebsiella* phage, therapeutic and prophylactic treatment. **(B)** Survival curves of *C. elegans* infected and treated with bacteria and phage ratio of 1:1 i.e. 10^5^ CFU/mL and 10^5^ PFU/mL, **(C)** Survival curves of *C. elegans* infected and treated with bacteria and phage ratio of 1:10 i.e. 10^5^ CFU/mL and 10^6^ PFU/mL, **(D)** Survival curves of *C. elegans* infected and treated with bacteria and phage ratio of 1:100 i.e. 10^5^ CFU/ mL and 10^7^ PFU/mL. The survival curves were plotted using Kaplan-Meier method and log-rank test was used to analyze the difference in survival rates in GraphPad Prism 7.0. A statistically significant difference (p<0.05) was observed in the treatment groups.

As a fourth pathogenic strain, we tested an *Enterobacter cloacae* isolate although *Enterobacter* is known to often be part of the commensal flora in *C. elegans* (37). However, no *Enterobacter* species were obtained in our *C. elegans* when attempting to specifically isolate such species. Similarly to the pathogenic *E. coli* strain 140 compared to the *E. coli* feeding strain OP50, the observations we made illustrates that strains of the same species can exhibit fundamentally different effects on the host, likely due to virulence factors. The *E. cloacae* strain 140 is a clinical isolate (from a urine sample) obtained from a diagnostic center in Chennai (India) that resulted in mortality of infected nematodes with a TD_mean_ of 3 days (Fig.5A). While purified *Enterobacter* phage myPSH1140 alone had no negative effect on the nematodes (survival >90%), the therapeutic treatment group showed increased survival to 85% at 5 days when 1:100 was used (Fig.5D). However both 1:1 and 1:10 had survival up to 75% at 5 days (Fig.5B,C). The prophylactic phage treatment allowed survival of 85% of the nematodes when 1:100 was used but up to 80% survival at the same time point (Fig.5B,C,D).

**Figure 5:**
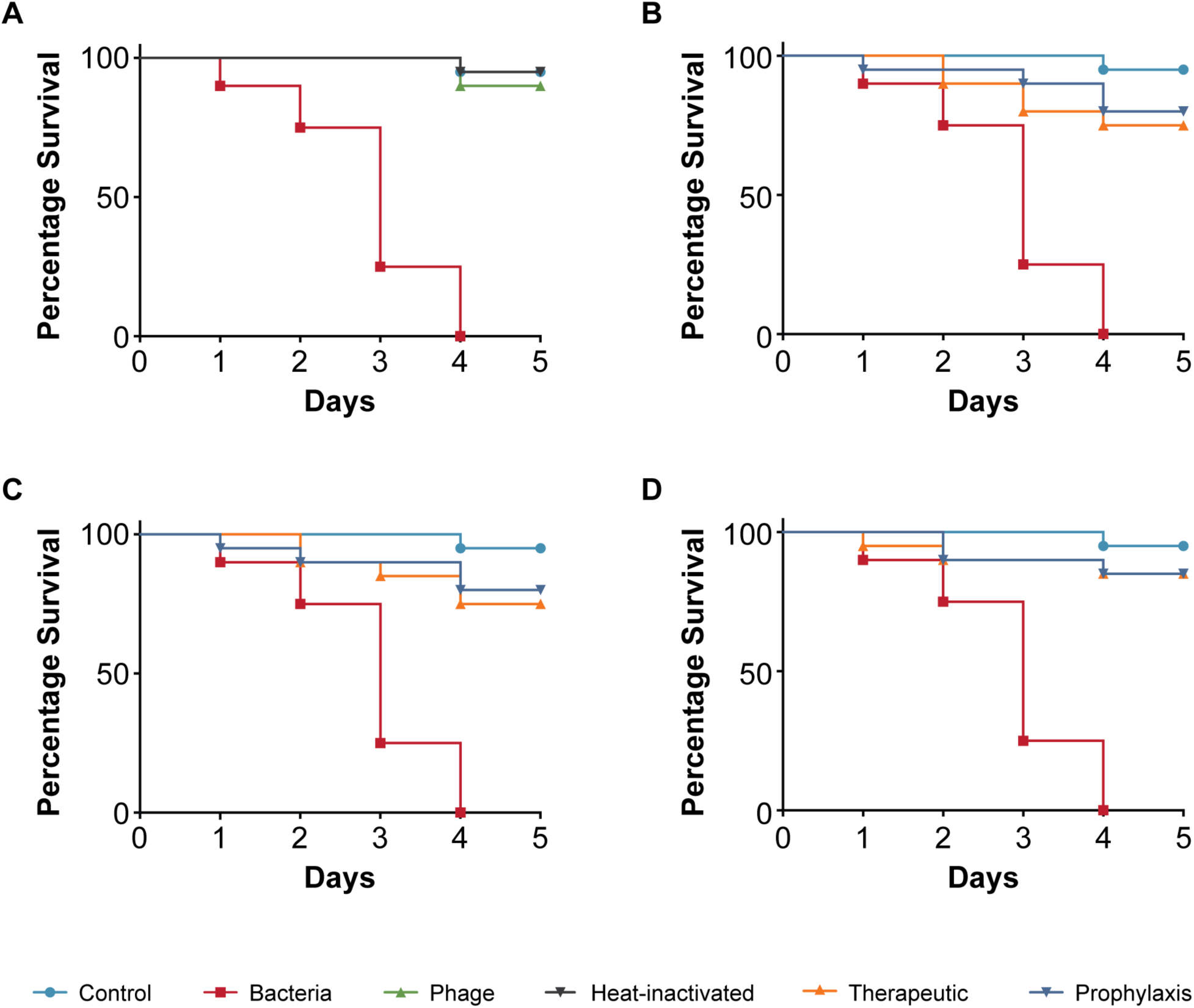
Pathogenicity of *Enterobacter cloacae* strain 140 in *C. elegans* and efficacy of *Enterobacter* phage myPSH1140 against *E. cloacae* infections. The control group consisted of *C. elegans* fed with *E. coli* OP50 and exposed to *E. cloacae* strain 140 (OD_600_=0.6) that kills *C. elegans* in liquid medium. 20 nematodes were used in each group. Representative survival curves of *C. elegans* following infection by *E. cloacae* strain 140 in **(A)** liquid medium consisting of M9 buffer and *E. cloacae* culture or *Enterobacter* phage or heat-inactivated bacteria and **(B,C,D)** survival curves of *C. elegans* following infection with *E. cloacae* strain 140 and treatment with *Enterobacter* phage, therapeutic and prophylactic treatment. **(B)** Survival curves of *C. elegans* infected and treated with bacteria and phage ratio of 1:1 i.e. 10^5^ CFU/mL and 10^5^ PFU/ mL, **(C)** Survival curves of *C. elegans* infected and treated with bacteria and phage ratio of 1:10 i.e. 10^5^ CFU/mL and 10^6^ PFU/mL, **(D)** Survival curves of *C. elegans* infected and treated with bacteria and phage ratio of 1:100 i.e. 10^5^ CFU/mL and 10^7^ PFU/mL. The survival curves were plotted using Kaplan-Meier method and log-rank test was used to analyze the difference in survival rates in GraphPad Prism 7.0. A statistically significant difference (p<0.05) was observed in the treatment groups.

Virulence is often assessed with a single pathogen. However, many infections in clinical practice are not caused by a single pathogen but by several, sometimes with opportunistic bacteria complicating the infection. Such poly-microbial infections are rarely investigated in the lab. Here we used a mixture of the pathogens (*E. coli* 131, *E. coli* 311, *K. pneumoniae* 235, *E. cloacae*) which we had tested individually and investigated them in our animal model. Such poly-microbial infections significantly reduced the survival percentage to 50% within 2 days, possibly displaying synergistic virulence effects, not cumulative ones as we used the same final number of cells (Fig.6A). When we next investigated if phages had a beneficial effect decreasing mortality of the nematodes, we prepared a so-called “phage cocktail” which contained phages infecting all the pathogens we used in our poly-microbial infection test. When investigating the effect of the phage cocktail alone on the nematodes, no negative influence on survival was observed. Using the mixture of phages in worms exposed to the pathogens, the survivability of nematodes was up to 75% (therapeutic) and 80% (prophylaxis) when the phage concentration was higher i.e. 1:100 (Fig.6D). A reduction in survival was observed when the phage concentration against bacteria was reduced (Fig.6B,C). This observation demonstrates the efficacy of phage cocktails in reducing the bacterial load caused by the tested poly-microbial infections (Fig.6B,C,D).

**Figure 6:**
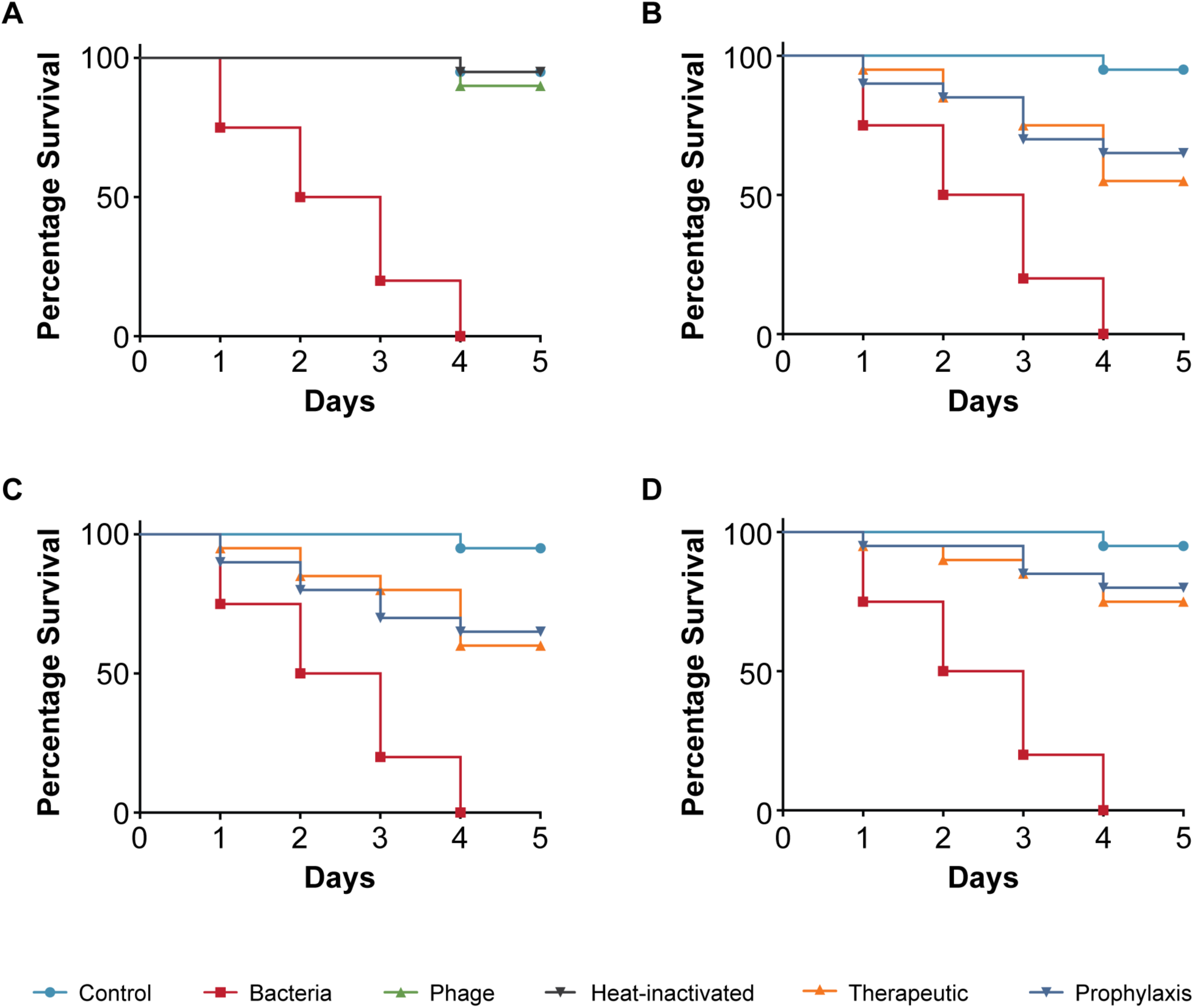
Pathogenicity of multi-bacterial culture in *C. elegans* and efficacy of phage cocktails against poly-microbial infections. The control group consisted of *C. elegans* fed with *E. coli* OP50. 20 nematodes were used in each group. Poly-microbial culture (*E. coli* 131, *E. coli* 311, *K. pneumoniae* 235, *E. cloacae* 140) kills *C. elegans* in liquid medium. Representative survival curves of *C. elegans* following infection by poly-microbial culture in **(A)** liquid medium consisting of M9 buffer and bacterial culture or phage cocktail or head-inactivated bacteria and **(B,C,D)** survival curves of *C. elegans* following infection with poly-microbial bacteria and treatment with phage cocktail, therapeutic and prophylactic treatment. **(B)** Survival curves of *C. elegans* infected and treated with poly-microbial bacteria and phage cocktail ratio of 1:1 i.e. 10^5^ CFU/mL and 10^5^ PFU/mL, **(C)** Survival curves of *C. elegans* infected and treated with poly-microbial bacteria and phage cocktail ratio of 1:10 i.e. 10^5^ CFU/mL and 10^6^ PFU/mL, **(D)** Survival curves of *C. elegans* infected and treated with poly-microbial bacteria and phage cocktail ratio of 1:100 i.e. 10^5^ CFU/mL and 10^7^ PFU/mL. Survival curves were plotted using Kaplan-Meier method and log-rank test was used to analyze the difference in survival rates in GraphPad Prism 7.0. A statistically significant difference (p<0.05) was observed in the treatment groups.

As phages can infect and inactivate bacteria in the media surrounding the worm, we next investigated if the phages are being taken up by the nematodes. The amount of phage particles inside the worms was reduced to 10^2^ PFU/mL from an initial concentration of 10^6^ PFU/mL after 4 days in a solution containing only the phage (Fig.7). However, a ten-fold higher concentration of phages (∼10^3^ PFU/mL) was found in the therapeutic and in the prophylactic treatment groups indicating that phage replication occur inside the nematode (Fig.7).

**Figure 7:**
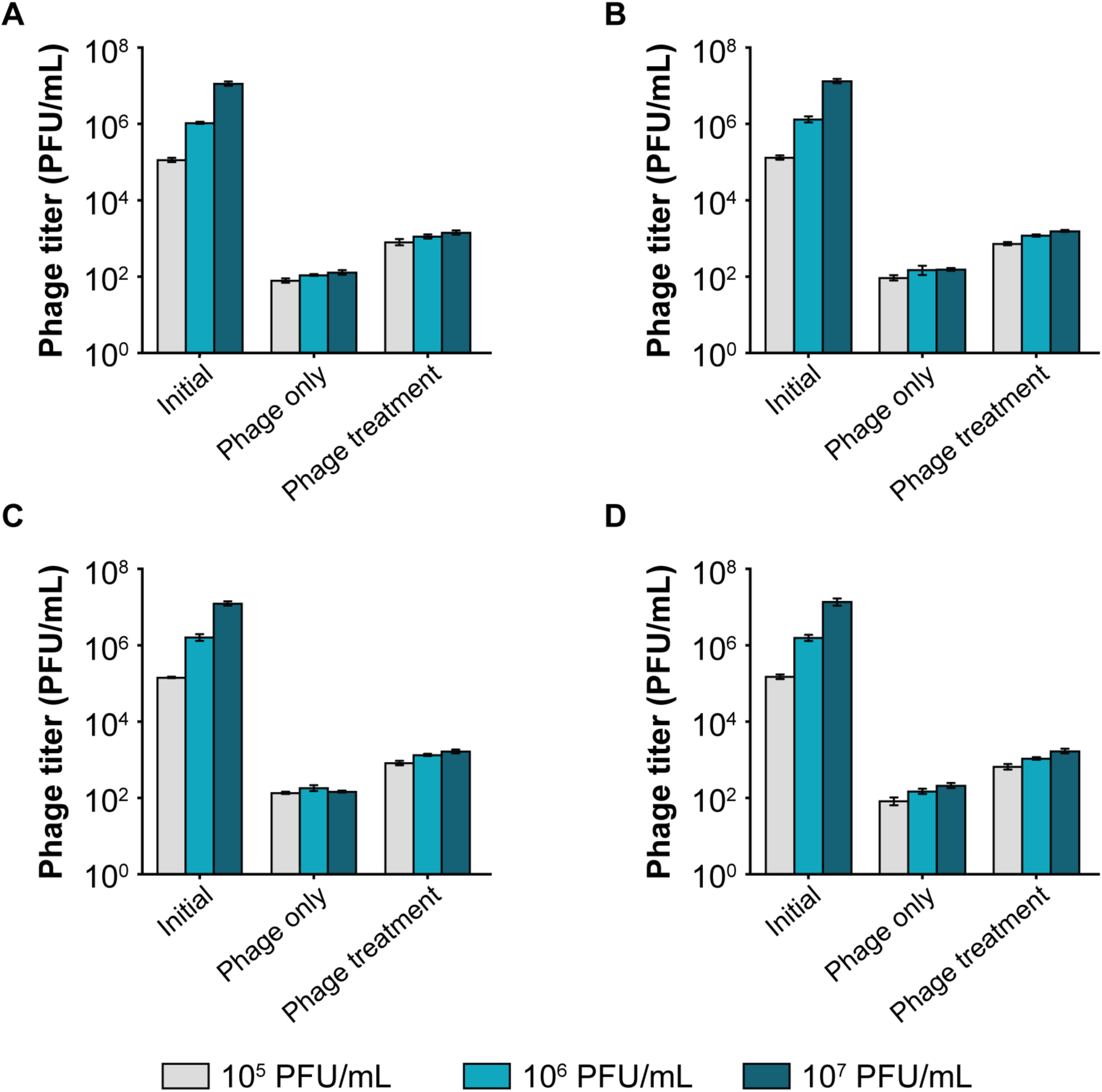
Bacteriophage enumeration from *C. elegans* after 4 days of exposure under two different conditions, (1) Phage only i.e. *C. elegans* was introduced to phage alone, and (2) Phage treatment i.e. *C. elegans* infected with bacteria was treated with phages. **(A)** *Escherichia* phage myPSH1131, **(B)** *Escherichia* phage myPSH2311, **(C)** *Klebsiella* phage myPSH1235 and **(D)** *Enterobacter* phage myPSH1140. After 4 days of treating *C. elegans* with phages, the nematodes were washed, grained and titrated for the presence of phages. The number of phages in treatment group was 10 times higher than the untreated group. The error bars represent the standard mean of three independent experiments.

For all four bacteriophages, the phage treatment resulted in a 6-fold or higher increase in nematode survival. In all cases, the prophylactic treatment resulted in a higher percentage of nematode survival compared to the therapeutic treatment. This is possibly due to a decrease in bacterial load as phages are able to kill bacteria that are not yet being ingested by the nematode. Regardless of this issue which is not straight forward to test, our work has clearly established that *C. elegans* can serve as a robust and reproducible, statistically valuable testing platform not only for assessing the virulence of bacterial pathogens but more importantly the efficacy of potentially therapeutic phages, a first step towards ensuring safe deployment in phage therapy.

## Discussion

We have successfully established a liquid-based *C. elegans* screening platform to investigate the efficacy of potential therapeutic bacteriophages. This robust and reproducible approach to study the antibacterial potential of lytic phages using a whole-animal infection model is valuable not only because of its ease but also due to better statistics with higher numbers of animals compared to other established assays. We were able to show that our method appears to be employable for a diverse range of pathogens which cause diseases in humans, as they also cause infections in *C. elegans* (38). Most studies involving *C. elegans* are routinely performed on agar plates. However, the use of a liquid-based screening method has several advantages including time efficiency and easy experimental design, as components can be added to the solution which allows for an even exposure of the nematodes to bacteria and/or bacteriophage. The screening on titer-plates allows quick nematode scoring i.e. makes easy and clear distinguishing of nematodes, as the colorless M9 buffer allows the nematodes to be observed clearly in contrast to solid media. Even though some experience is required to conduct worm survival counts, the skill can be acquired easily. A liquid-based screening with *C. elegans w*as previously found to be suitable to address the virulence of *Staphylococcus aureus* (25). The method employs an approach similar to ours, using *C. elegans* as a model organism in liquid media requires high-end equipment and the use of bacteriophages has not been established with this method (26). In addition, our assay is easier to use, faster and less cost-intensive. However, in its current form, our assay can be regarded as semi-quantitative as a human operator evaluates the viability of the nematodes. Thus the development of a fully quantitative assay would be an improvement for future optimisation of the method. Potential approaches might include a robotic, automated counting of worms using e.g. specifically developed image processing software (39), or an antibody-based ELISA against life/ dead specific marker proteins.

Using the simple animal model in our assay, we have also demonstrated the efficacy of four bacteriophages in increasing the survival of *C. elegans* infected with bacterial pathogens. The four bacteriophages (*Escherichia* phage myPSH1131, *Escherichia* phage myPSH2311, *Klebsiella* phage myPSH1235 and *Enterobacter* phage myPSH1140) were found to be effective in eliminating the bacterial load, increasing the lifespan of *C. elegans* (Fig.2-5). Although the higher concentration of bacteria-phage ratio (1:100 or 10^5^ CFU/mL: 10^7^ PFU/mL) showed up to 90% nematode survival, the lesser concentrations of 1:1 and 1:10 had at least 80% survival. Therefore, survivability of the nematodes was observed irrespective of the phage concentration. Also in combination, as a phage cocktail, the phages displayed highly beneficial effects in limiting the impact of poly-microbial infections on the nematodes. To the best of our knowledge, this is the first study to report the successful use of *C. elegans* to test the efficacy of phages in a liquid assay format.

The data from our assay confirms results we have previously obtained in experiments using waxworms (15). In our previous study, when waxworms were infected with bacteria (10^8^ CFU/mL) and treated with phages (10^4^ PFU/mL), up to 80% larval survival was noted (15). In this study, perhaps not surprisingly, the prophylactic treatment allows for better nematode survival rates than the therapeutic treatment where an infection is first established before treating the worm with phages. However, even in the treatment regime we obtained statistically significant high survival rates which demonstrate that bacteriophages are able to protect or cure *C. elegans* from bacterial infections caused by the four different clinical strains, also when combined in an effort to reproduce a ploy-microbial infection. We believe that this is the first study to test the pathogenesis of poly-microbial infections in *C. elegans* and also to use phage cocktails against the poly-microbial infections.

As phage therapy becomes an increasingly used clinical intervention to treat MDR infections, the use of simple live animal models in robust and reliable assays like the one we have established for *C. elegans* will facilitate the therapeutic evaluation of phages. While innate immune factors are conserved among nematodes, insects, and mammals, there are distinct differences among lower animals (40 - new reference here: 10.1016/j.molimm.2007.09.030). Both, *Galleria* and *Caenorhabditis* lack an adaptive immune system, however studies on the innate immune systems of insects and nematodes have indicated that there are differences in pattern- recognition- molecules and signalling factors. Although results of our work do not indicate any difference (such as a higher/lower tolerance to certain pathogens in *C. elegans*), the insect and the nematode model might show variances when evaluating the virulence of bacteria. However, for clinical phage therapy, in particular for human applications, an animal model with an adaptive immune system should be used. Thus an invertebrate-based screening can only serve as the first stage to evaluate the therapeutic potential of a phage, yet in our case as a rapid and inexpensive one. Such robust, reliable and potentially high-throughput methods can be considered one of the prerequisites for the implementation of phage therapy, from the discovery of environmental phages or the construction of synthetic viruses, to the phage solution ready for medical treatment.

## Conclusion

The data provided in this study demonstrates the ability of a liquid-based assay to test the antimicrobial efficacy of bacteriophages to increase the nematode survival when infected with bacterial pathogens. This allows the activity of phages to be tested before large scale pre-clinical studies in mouse models. While the use of *C. elegans* was explored in this study by manual operation, observation and analysis, modifications to our test system could allow the adaptation to establish a high-throughput screening platform for hundreds of bacteriophages in parallel.

## Materials and methods

### Bacterial strains and bacteriophages

A total of four clinical bacterial strains, *Escherichia coli* 131 (Enteropathogenic), *Escherichia coli* 311 (Enterohemorrhagic), *Klebsiella pneumoniae* 235 and *Enterobacter cloacae* 140 were previously studied and chosen for the study (41,42). *E. coli* pathotypes were identified by polymerase chain reaction as detailed in our previous studies (41,42). Bacterial strains including the non-pathogenic “nematode feeding bacterium” *E. coli* OP50 were grown on Lysogeny Broth (LB) agar at 37°C and stored in LB medium. For the assay, bacterial strains were grown overnight at 37°C with shaking at 120 rpm, then the culture was diluted to OD_600_=0.6 (∼10^5^ CFU/mL) unless stated otherwise. Heat-inactivated bacteria was prepared by incubating the cells at 65°C for 40 min (43). 1 mL of cells was pelleted by centrifugation at 4000 × g for 10 minutes before washing the pellet thrice with PBS and finally resuspending it in 1 mL PBS. Four bacteriophages, *Escherichia* phage myPSH1131, *Escherichia* phage myPSH2311, *Klebsiella* phage myPSH1235 and *Enterobacter* phage myPSH1140 were used in this study (41,42). The purified bacteriophages were prepared at 10^5^ – 10^7^ PFU/mL as described previously (41,42). All the bacterial strains and bacteriophages used in this study are available at Antibiotic Resistance and Phage Therapy Laboratory, Vellore Institute of Technology, Vellore, India.

### C. elegans maintenance

Bristol N2 (wild-type) *C. elegans* was used in this study and was kindly provided by Dr. N. Elangovan, Periyar University, Salem, India. The nematodes were maintained and propagated on nematode growth media (NGM; 17 g agar, 3 g NaCl, 2.5 g peptone, 0.5 mL of 1M CaCl_2_, 1 mL of 5 mg/mL cholesterol, 1 mL of 1M MgSO_4_, 25mL KH_2_PO_4_ buffer [pH 6.0] per litre) plates that carries *E. coli* OP50 as a source of food at 20°C by standard protocols (44). The adult worms were exposed to 5 M sodium hydroxide and 5% bleach to collect eggs that were then incubated in M9 medium (6 g Na_2_HPO_4_, 3 g KH_2_PO_4_, 5 g NaCl, 0.25 g MgSO_4_.7H_2_O). Worms take approximately 12 hours to hatch, and only mature eggs allow the development of viable nematodes by which synchronization was archived to obtain worms of the same age (45). For the experiments, age synchronized L4 worms were incubated in the presence of tryptic soy broth (TSB), *E. coli*, *K. pneumoniae*, *E. cloacae* and/or bacteriophage of *Escherichia* phage myPSH1131, *Escherichia* phage myPSH2311, *Klebsiella* phage myPSH1235 and *Enterobacter* phage myPSH1140 in 96 well microtitre plates containing 100 µL of M9 buffer in each well.

### Testing phage efficacy in C. elegans model

Two types of assays were performed i.e. therapeutic treatment and prophylactic treatment. For the liquid-based assay, a 96-well microtiter plate was filled with M9 buffer to which an overnight culture of *E. coli* OP50 (feeding bacteria) or the equivalent amount of pathogenic bacteria, with or without phage was added (Fig.8 ). Then 20 mature L4 nematodes were transferred into the solution. The total volume in the microtiter plate well was maintained at 100 µL. In order to test the robustness of the experiments, different concentrations of pathogenic bacteria was used to test the infectivity. Accordingly, overnight grown bacterial cultures were diluted to 10^3^, 10^5^ and 10^7^ CFU/mL for testing. The efficacy of phage treatment was studied using varying concentrations of bacteria and phage i.e. 1:1, 1:10 and 1:100 which were 10^5^ CFU/mL: 10^5^ PFU/mL and 10^5^ CFU/mL: 10^6^ PFU/mL and 10^5^ CFU/mL: 10^7^ PFU/mL.

**Figure 8:**
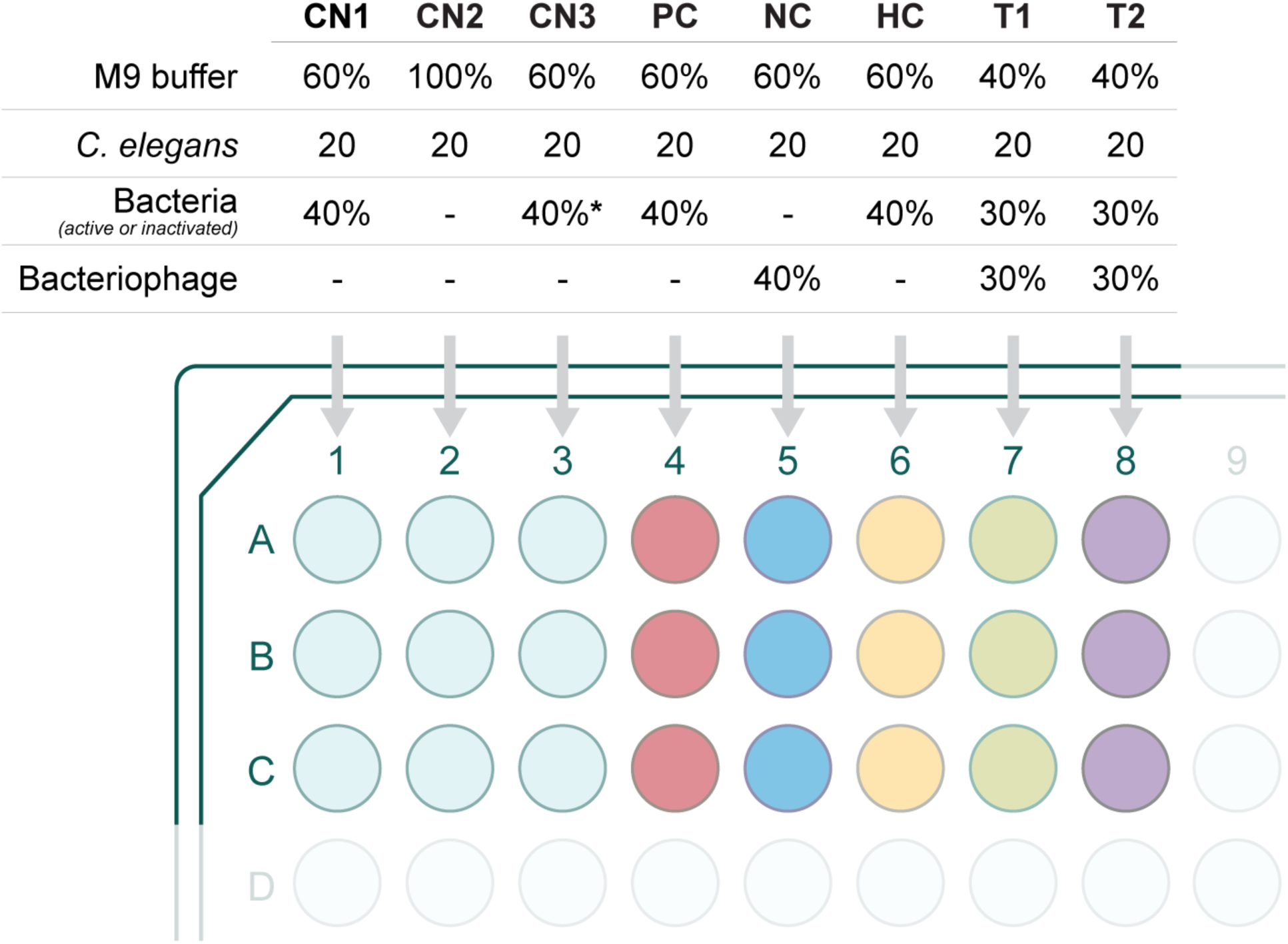
Representation of the study groups used to evaluate the efficacy of bacteriophages in treating bacterial infections in *C. elegans* model. In a 96 well microtiter plate, experiments were conducted in triplicate. Details are as follows: Negative controls **CN1** - M9 buffer (60%), *E. coli* OP50 (40%) and 20 nematodes; **CN2** - M9 buffer and 20 nematodes; **CN3** - M9 buffer (60%), TSB (40%) and 20 nematodes. **PC** (infection control) - M9 buffer (60%), overnight bacterial culture of *E. coli* or *K. pneumoniae* or *E. cloacae* (40%) and 20 nematodes. **NC** (phage control) - M9 buffer (60%), bacteriophage (*Escherichia* phage myPSH1131, *Escherichia* phage myPSH2311, *Klebsiella* phage myPSH1235 and *Enterobacter* phage myPSH1140) (40%) and 20 nematodes. **HC** (heat-inactivated bacteria control) - M9 buffer (60%), heat-killed bacterial culture of *E. coli* or *K. pneumoniae* or *E. cloacae* (40%) and 20 nematodes. **T1** (treatment group, phages were added 2 hrs after bacterial infection) - M9 buffer (40%), overnight bacterial culture of *E. coli* or *K. pneumoniae* or *E. cloacae* (30%), bacteriophage (30%) and 20 nematodes. **T2** (prophylactic treatment group, phages were added 1 hr before the bacterial infection) - M9 buffer (40%), bacteriophage (30%), overnight bacterial culture of *E. coli* or *K. pneumoniae* or *E. cloacae* (30%) and 20 nematodes. * indicates TSB was used instead of bacteria.

**Group 1** (control) consisted of M9 buffer (60%) with *E. coli* OP50, with ∼10^5^ CFU/mL (40%) and 20 nematodes. **Group 2** M9 buffer, 20 nematodes, no bacteria. **Group 3** M9 buffer (60%), TSB (40%), 20 nematodes. Both groups 2 and 3 were used as experimental controls. **Group 4** (infection control): M9 buffer (60%), bacterial pathogen (*E. coli* or *K. pneumoniae* or *E. cloacae*, 40%), 20 nematodes. **Group 5** (heat inactivated bacteria): M9 buffer (60%), heat-inactivated bacteria (*E. coli* or *K. pneumoniae* or *E. cloacae*, 40%), 20 nematodes. **Group 6** (phage toxicity test): M9 buffer (60%), bacteriophage (*Escherichia* phage myPSH1131, *Escherichia* phage myPSH2311, *Klebsiella* phage myPSH1235 and *Enterobacter* phage myPSH1140, 40%) and 20 nematodes. **Group 7** (therapeutic treatment group): M9 buffer (40%), bacterial pathogen (*E. coli* or *K. pneumoniae* or *E. cloacae,* 30%), 20 nematodes; bacteriophage (30%) were added after 2 hours of exposure to bacteria. **Group 8** (prophylactic treatment group): M9 buffer (40%), bacteriophage (30%), 20 nematodes; bacterial cultures of *E. coli* or *K. pneumoniae* or *E. cloacae* (30%) were added after 1 hour. Plates were incubated at 20°C and survival of the nematodes was monitored every 24 hours for 5 days. The results were evaluated based on live nematodes (moving) and dead nematodes (lack of movement), see Fig.1. The L4 stage nematodes were chosen for the study because even though the lifespan of *C. elegans* is 20 ± 3 days, nematodes drastically reduce motility when ageing, making it difficult to establish if infected (46). All the experiments were repeated a minimum of three times for statistical significance.

The efficacy of a phage cocktail was evaluated by establishing poly-microbial infections. To this end, the bacterial cultures were mixed at equal volumes containing the same CFU numbers. Heat-inactivated bacteria were also prepared from the mixed culture. The phage cocktail was prepared by mixing the bacteriophages at equal volumes with identical PFUs. The efficacy of phage treatment was studied using varying concentrations of bacteria and phage i.e. 1:1, 1:10 and 1:100 which were 10^5^ CFU/mL: 10^5^ PFU/mL, 10^5^ CFU/mL: 10^6^ PFU/mL and 10^5^ CFU/mL: 10^7^ PFU/mL. **Group 1** consisted of M9 buffer (60%) with *E. coli* OP50 (40%) and 20 nematodes. **Group 2** M9 buffer, 20 nematodes, no bacteria. **Group 3** M9 buffer (60%), TSB (40%), 20 nematodes. **Group 4** (infection control): M9 buffer (60%), mixed bacterial culture (*E. coli*, *K. pneumoniae*, *E. cloacae*, 40%), 20 nematodes. **Group 5** (heat inactivated bacteria): M9 buffer (60%), heat-killed bacteria (poly-microbial), 20 nematodes. **Group 6** (phage toxicity test): M9 buffer (60%), phage cocktail (consists of four phages, 40%) and 20 nematodes. **Group 7** (therapeutic treatment group): M9 buffer (40%), mixed bacterial culture (*E. coli*, *K. pneumoniae*, *E. cloacae*, 40%), 20 nematodes; phage cocktail (30%) was added after 2 hours of exposure to the bacteria. **Group 8** (prophylactic treatment group): M9 buffer (40%), phage cocktail (30%), 20 nematodes; the poly-microbial culture containing *E. coli*, *K. pneumoniae*, *E. cloacae* (30%) was added after 1 hour. Plates were incubated at 20°C and survival of the nematodes was monitored every 24 hours for 5 days. The results were evaluated and statistical analysis was performed.

### Enumeration of bacteriophages from the treated nematodes

To analyse the presence of bacteriophages inside the nematodes or uptake of bacteriophages by the nematodes, phages numbers were determined as follows: Briefly, after 4 days, 10 nematodes were removed from groups 5, 6 and 7 which are phage only, therapeutic treatment and prophylactic treatment, respectively, to determine the phage titer using double agar overlay method. In brief, the nematodes were washed thrice with M9 buffer, vortexed, ground (with mortar and pestle) and centrifuged at 10,000 × g for 5 min. The supernatant was used to determine the phage titer. All bacteriophages were used alone (not in combination) and the experiments were repeated three times for statistical significance.

### Statistical analysis

Survival curves were plotted using Kaplan-Meier method and *log-rank test* was used to calculate the difference in survival rates using GraphPad Prism software 7.0 (GraphPad Software, Inc., La Jolla, USA). P<0.05 was considered as statistically significant (*log-rank test*).

## Acknowledgement

The authors would like to thank Vellore Institute of Technology for providing ‘VIT Seed Grant’ and the support provided by the Zhejiang Province Postdoctoral Research Fund (ZJ2020151) to Prasanth Manohar. The authors gratefully acknowledge Caenorhabditis Genetic Centre (CGC, University of Minnesota, MN, USA) which is funded by NIH Office of Research Infrastructure Programs (P40 OD010440) for providing Bristol N2 (wild type) used in this work.

## Supplementary figures

**S. figure 1:**
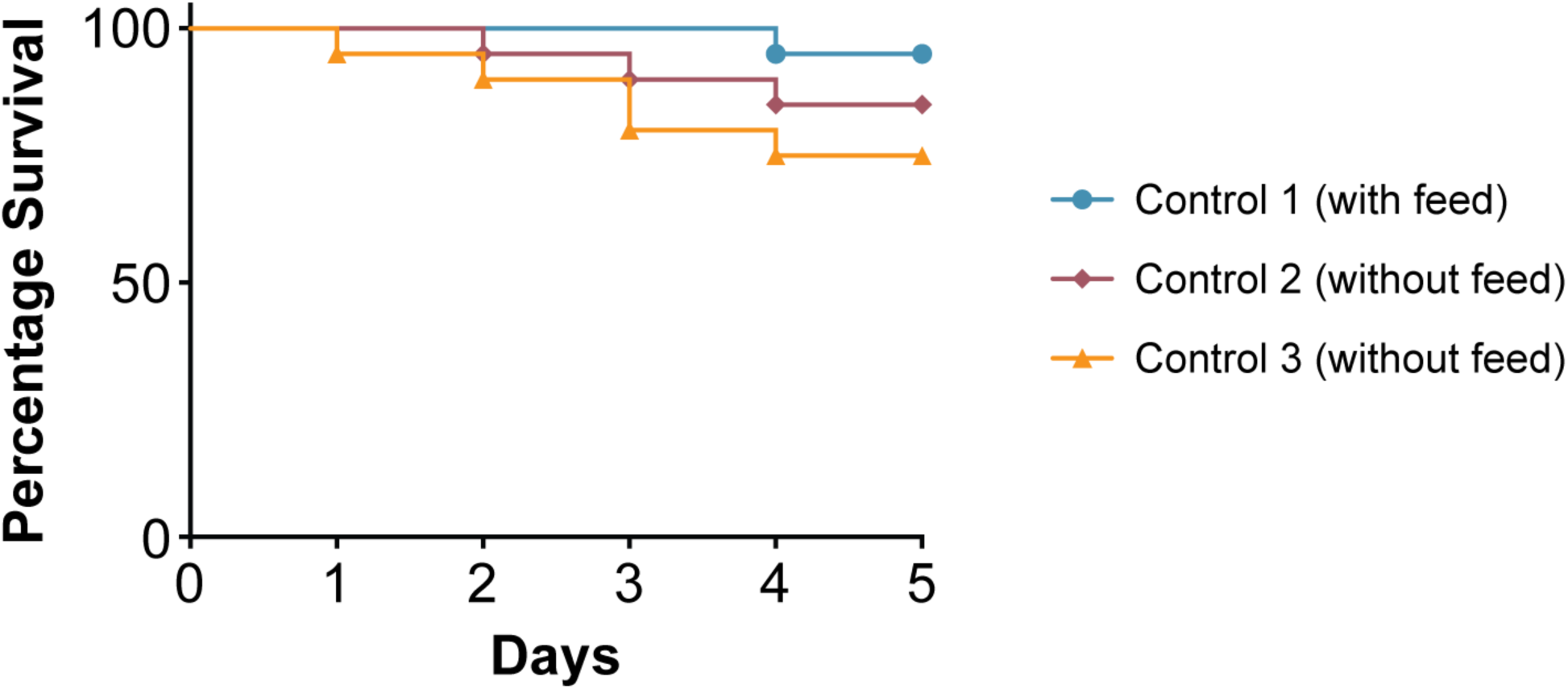
Survival curves of nematodes (*C. elegans*). Nematodes were kept in varying growth conditions; Control 1: nematodes were fed with *E. coli* OP50, Control 2 (liquid media) & Control 3 (solid-media): nematodes were not fed. Survival curves were plotted using Kaplan-Meier method and log-rank test was used to analyze the difference in survival rates in GraphPad Prism 7.0.

**S. figure 2:**
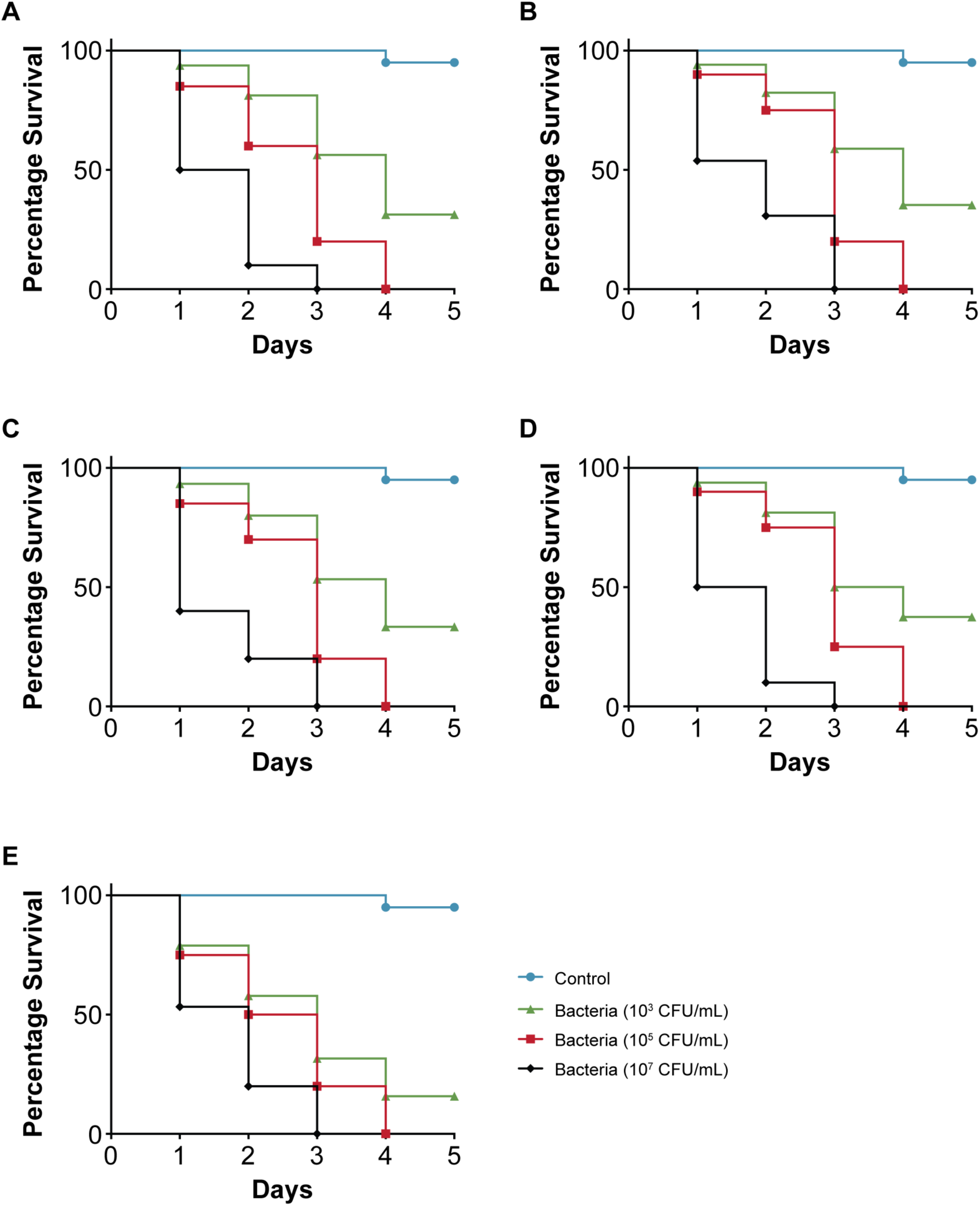
Survival curves of nematodes (*C. elegans*) when infected with pathogenic bacteria at different concentrations, 10^3^, 10^5^ and 10^7^ CFU/mL. **(A)** *C. elegans* infected with *Escherichia coli* 131; **(B)** *C. elegans* infected with *Escherichia coli* 311; **(C)** *C. elegans* infected with *Klebsiella pneumoniae* 235; **(D)** *C. elegans* infected with *Enterobacter cloacae* 140. The survival curves were plotted using Kaplan-Meier method in GraphPad Prism 7.0.

**S. figure 3:**
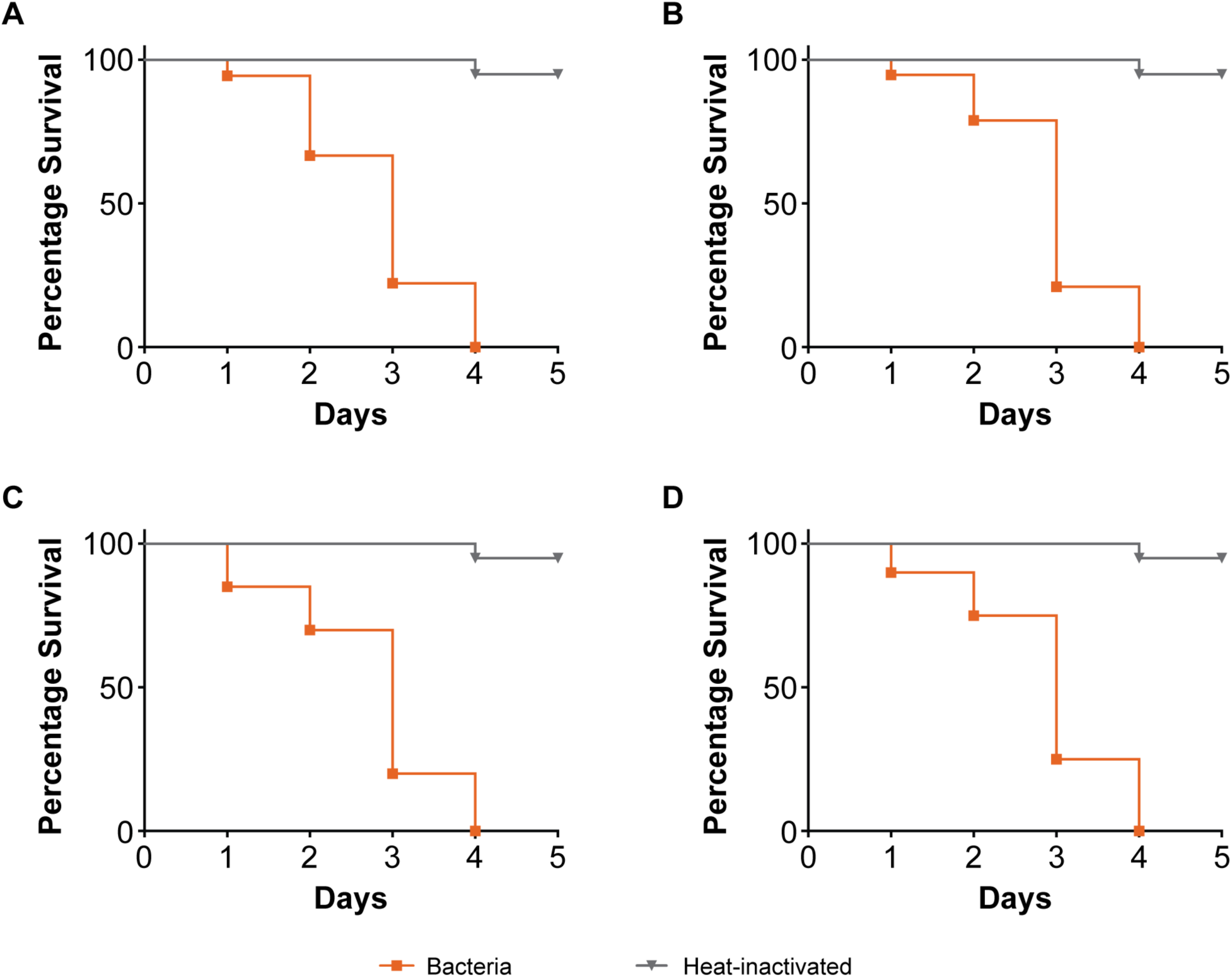
Survival curves of nematodes (*C. elegans*) when infected with live bacteria and heat-killed bacteria. **(A)** *C. elegans* infected with *Escherichia coli* 131; **(B)** *C. elegans* infected with *Escherichia coli* 311; **(C)** *C. elegans* infected with *Klebsiella pneumoniae* 235; **(D)** *C. elegans* infected with *Enterobacter cloacae* 140. The survival curves were plotted using Kaplan-Meier method in GraphPad Prism 7.0.

